# Analysis of the transcriptional logic governing differential spatial expression in Hh target genes

**DOI:** 10.1101/333260

**Authors:** Manuel Cambón, Óscar Sánchez

## Abstract

This work provides theoretical tools to analyse the transcriptional effects of certain biochemical mechanisms (i.e. affinity and cooperativity) that have been proposed in previous literature to explain the differential spatial expression of Hedgehog target genes involved in *Drosophila* development. Specifically we have focused on the expression of *decapentaplegic* and *patched*. The transcription of these genes is believed to be controlled by opposing gradients of the activator and repressor forms of the transcription factor Cubitus interruptus (Ci). This study is based on a thermodynamic approach, which provides expression rates for these genes. These expression rates are controlled by transcription factors which are competing and cooperating for common binding sites. We have made mathematical representations of the different expression rates which depend on multiple factors and variables. The expressions obtained with the model have been refined to produce simpler equivalent formulae which allow for their mathematical analysis. Thanks to this, we can evaluate the correlation between the different interactions involved in transcription and the biological features observed at tissular level. These mathematical models can be applied to other morphogenes to help understand the complex transcriptional logic of opposing activator and repressor gradients.

**Author summary:** Morphogenic differentiation is a complex process that involves emission, reception and cellular response to different signals. It is well known that the same morphogenic signal can give rise to different cellular transcriptional responses that usually depend, among other factors, on transcription factors. In concordance with the activator threshold model, classically it has been distinguished between high and low threshold target genes in order to explain how cells receiving the same signal can activate different genes. However, in particular cases where the transcription is controlled by two opposing transcription factors, it has been tested that this logic is not valid. This motivates the necessity for describing new theoretical models in order to understand better these cellular responses. By a theoretical analysis we have deduced different versions of transcriptional logic that are significantly determined by how the opposing transcription factors cooperate between them in the transcription process. We have also tested these different scenarios focussing on the Drosophila Hh target genes, and we have reproduced similar conclusions to the ones obtained by other methodologies.

## Introduction

Hedgehog (Hh) is a morphogen, a signalling protein that induces several cellular responses. It is involved in the development of different biological systems, for example that of, the Drosophila melanogaster fly. In *Drosophila*’s wing imaginal disc the secretion of Hh from the Posterior compartment cells induces the expression of several target genes inside the cells in the Anterior compartment. Among them are *decapentaplegic (dpp*) and *patched* (*ptc*). Both give rise to the synthesis of their corresponding proteins, Dpp and Ptc, which are essential for the wing central domain development [20,21].

However, it is known that the same signal of Hh produces different spatial expression of theses genes. That is to say, the expression of *ptc* is only limited to disc zones close to the *Anterior/Posterior* (A/P) border with high Hh concentrations, while *dpp* expresses in a broader disc range under low Hh concentrations. This poses a question: Why does the same signal give rise to different spatial expressions for different genes: The answer to this question is still under debate. The current understanding is that both genes respond, basically, to the same principles that we list below.

Hh transcriptionally controls both Dpp and Ptc through the *transcription factors* (TFs) Cubitus interruptus (Ci). It dictates the activity of RNA polymerase enzymes (RNAP), which controls the genetic transcription via the synthesis of Ribonucleic Acid (mRNA). This process requires the binding of RNAP to some specific sites on the DNA chain called promoters. However, the transcription rate of the target genes not only depends on the total concentration of RNAP in the system, but also is controlled by the protein Ci. Ci is present in two opposite forms: activator and repressor. The activators, CiA, attempt to promote the transcription rate while the repressors, CiR, attempt to decrease it. Hh signal affects the balance between both forms of Ci, i.e., in the absence of Hh Ci appears in its repressed form but when Hh is absorbed by the cell, Cubitus changes it role presenting its activator form. So, the Hh gradient in the Anterior compartment creates opposing activator (CiA) and repressor (CiR) gradients. Furthermore both Ci forms need to bind specific DNA sites called enhancers or cis-regulatory sites which are different from the RNAP binding sites.

Some recent works [13,16,23] postulate that the reason for the differential spatial expression of these genes could be found in certain biochemical factors involved in the transcription process. Firstly, the binding of both RNAP and Ci in the promoter and enhancers is carried out by chemical reactions. These require some free energy that is commonly characterised by a magnitude called binding affinity. This affinity depends on several characteristics of the promoters and enhancers of each transcribed gene. In fact, in [16] it was observed that the enhancers with lower relative affinity seem to be necessary to obtain normal expression of *dpp* in regions of low signal. Secondly, it is possible that transcription factors that are already bound in some enhancers can modify the affinity of other binding elements. In this case, bound TFs may modify the free energy of a later binding reaction of either TF or RNAP. This process is generally termed cooperativity, however this can be positive or negative. If it facilitates the binding it is called (normal) cooperativity and if it impedes it, is called anti-cooperativity. In [14,15] it was proposed that the activator/repressor TFs modify the transcription rate by promoting or blocking respectively the recruitment of RNA polymerase. This implies that cooperativity or anti-cooperativity with the RNAP changes the promoter binding affinity. Thirdly, it is important to remark that, in the case of *Drosophila*, both CiA and CiR are in constant competition for the binding of a set of 3 enhancers. The combination of all these biochemical factors (competition, cooperativity and binding affinities) gives rise to a very complex balance between the concentration of activators and repressors making it difficult to discern their interacting effects at tissular level.

In [13], the spatial expression of both genes was related to the relatively higher affinity between Cubitus proteins and *dpp* enhancers, than the affinity between Cubitus proteins and *ptc* enhancers. Specifically, they confirm that, under moderate Hh signal, low-affinity sites produce activation, whereas high-affinity sites produce repression. In order to discriminate between the mechanisms that could give rise to a differential spatial expression, they contrasted the experimental observations with fittings to a thermodynamical model based on the ideas of Shea, Ackers and coworkers [2,19]. Furthermore, by fitting a repressor cooperativity model in [13] they observed that CiR plays a substantial role in the response to a moderate Hh signal. In [23] the authors also proposed that the cooperativity between repressors may play an important role in the change of the genetic expression along the imaginal disc, by using a mathematical model of occupancy competition between repressors and activators.

The large amount of biochemical variables that are present in the system calls for mathematical models that can shed some light on the origins of the differential spatial expression in the target genes of Hh, among others. The thermodynamic model proposed by Shea, Ackers and coworkers [2,19], also known as BEWARE [9] (Binding Equilibrium Weighted Average Rate Expression), is a method frequently used in the mathematical modelling of genetic transcription processes. See [3] or [8] for a general discussion/comparison with other modelling approaches as for instance Boolean models. However, this model gives rise to long and complex mathematical expressions even when there are only a few transcription factors involved. For the analysis of independent and specific binding sites and the analysis of two non competitive transcription factors, only simple mathematical expressions have previously been proposed [4,7]. It is difficult to decipher the biological effects in the model even if they are supported by numerical tools [24] because the expressions inherently involve a great number of constants and variables.

In this work we try to have a better understanding of the transcriptional logic of Hh target genes from a theoretical point of view by using a thermodynamic model. Our analysis proposes that the transcriptional logic in the presence of opposing activator/repressor gradients can exhibit different versions depending on the cooperativity between the transcription factors. In fact, the theoretical methodology we developed is able to demonstrate how the combination of similar biochemical factors applied under different frameworks gives rise to completely different transcriptional effects. We conclude that differential expression of Hh target genes is due to the combination of the differences in affinities between *dpp* and *ptc* binding sites and the cooperativity between repressors. Confirming the results obtained in [23]. However, this does not exclude that a different version of the transcriptional logic could be adopted by some other biological systems.

In the case of a single activator gradient our analysis represents the well accepted transcriptional logic which comes from the *activator threshold model*. This model explains the role of certain biochemical factors involved in the signalling interpretation. For example, differential affinities of activators for DNA elements [17] and cooperativity between activators [10,13]. High-affinity binding sites and cooperativity between activators benefit the binding of the activators to the enhancers, allowing the expression of genes at low activator concentrations, so here we observe a broader response within the activator gradient. In contrast, low-affinity sites and the absence of cooperativity between activators restrict the gene expression to high activator concentration regions. Although this rationale is well accepted in the single gradient scenario, it has not been succesfully applied in combinatorial interactions as, for instance in opposing activator-repressor gradients such as those we have described in Hh signalling [13,17]. In this case the balance between both gradients causes the existence of ranges of net activated/repressed cells, i.e., cells along the tissue that express higher/lower levels than basal level [13], and hence there is no global activation/repression. The *cellular expression ranges* (CERs) will not be determined only by signal intensity but also by net activated/repressed cellular ranges.

Our analysis suggests that biochemical differences between genes can affect both, signal modulation and changes in the net activated cellular ranges. That is, the variation of CERs can be explained by analysing the combination of both aspects. Another very interesting aspect is that using the same biochemical characteristics with different cooperativity between activators and repressors will give drastically different expression rates. Different types of cooperative interactions between transcription factors will produce variations in the transcription logic. Another very interesting aspect in our analysis is how the number of enhancers could modify the gene expression. In [13] a transgenic fly line carrying a GFP reporter with a single high-affinity Ci site was constructed in order to detect whether cooperativity between TFs played a important role in the signalling process. Even in absence of cooperativity, this variation of the number of enhancers could modify the gene expression, at least theoretically. This aspect, will play a key role in our argument in order to understand Hh target genes. Although they seem to play a central role, affinity and cooperativity between TFs are not the only biochemical factors involved in the interpretation of general morphogen signalling. In the development of the chick embryo neural tube, another paradigmatic morphogenetic patterning example, cells are differentiated in response to Sonic Hedgehog (Shh) morphogenetic signals [22]. In this case, the Shh signal balances the concentration of different versions of activators and repressors of the Gli family. These Gli TFs recognise target sequences which are very similar, however it has been proposed that some TFs have more potency as activators or repressors [10] than others. Thus, the different potency as activators and repressors could be another factor to be taken into account by using differential TFs-RNAPs cooperativity.

## Results

In this work we will study which of the previous factors can vary cellular expression ranges by analysing their combined effect on both, the modulation of the signal and the variation of the ranges of net activated/repressed cells. We begin with a modelling exercise that will provide the mathematical representation (operators) of expression levels to be later analysed. Firstly we apply the BEWARE method in order to obtain expression rates of a gene controlled by two opposing general transcription factors: the activator CiA and the repressor CiR. The “BEWARE operators” obtained include, for each gen:

- the transcription factors competing in order to bind one of *n* common cis-regulatory sites,
- separate binding affinities of the TFs depending on their activator or repressor form,
- cooperativity interactions between TFs and cooperativity/anti-cooperativity between TFs and RNAPs. We consider different process of TF-TF cooperativity that give rise to different expressions of the BEWARE operator:
  – non-cooperativity,
  – total cooperativity, when a bound TF modifies the binding affinity of any other transcription factor,
  – partial cooperativity, that only takes place between TFs that are of the same form, activators only interact with activators and repressors only interact with repressors.

In Fig. 1 (A) the upper scheme shows all these interactions with *n* = 3 binding sites. We have simplified the mathematical expressions of the BEWARE operators because in their original form it would be impossible to make the subsequent mathematical analysis.

**Fig 1.**
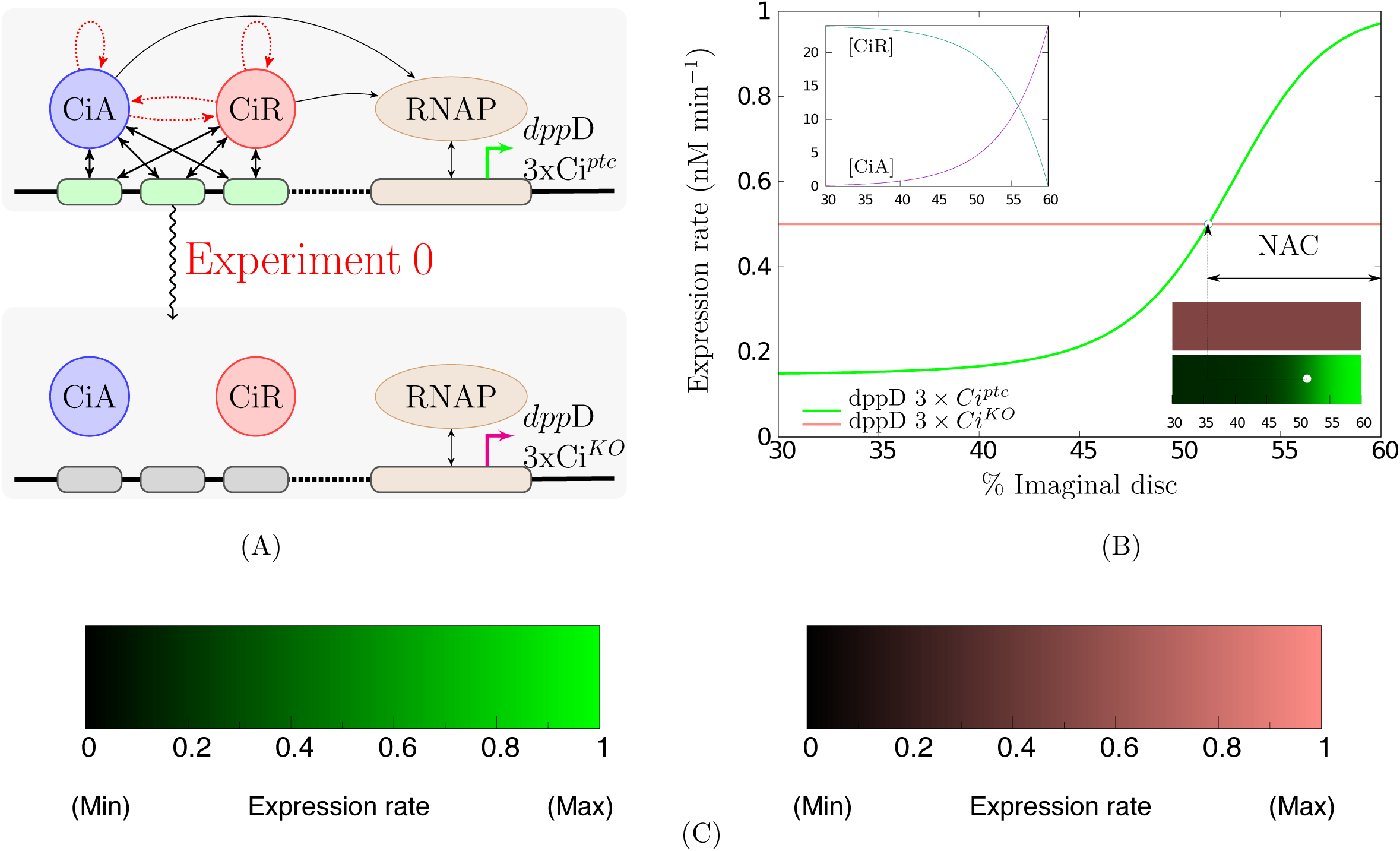
Net activated cellular (NAC) range described by a thermodynamic model. **A)** Schematic of the experiment for NAC range determination. The arrows represent all the possible interactions captured by a thermodynamic model determining the transcription rates: double-headed straight arrows show protein-DNA binding site affinities while single-headed black and red arrows are TFs-RNAP and TFs-TFs cooperativities respectively. The net activated cellular range of *dpp*D3xCi^*ptc*^, a reporter gene with a version of the *dpp* enhancer with three high-affinity binding sites, is obtained by comparing its theoretical transcriptional activity with the activity of *dpp*D3xCi^*KO*^, a gen containing different version of the *dpp* enhancer containing three null-affinity sites. Both cases are represented in the upper and lower schemes respectively. TFs binding sites are represented by rounded rectangles filled in green (high-affinity) or black (null affinity). **B)** Theoretical transcription rates predicted for both genes in cells of the Anterior compartment. This compartment occupies the 60% of the Drosophila imaginal disc and the Posterior compartment the rest (60% to 100%). The expression levels given by the BEWARE operators are between *0nM/min* and *1nM/min* being the basal level equal to 0.5nM/min. These reference expression levels have been chosen for a proper appreciation of signal modulation. Since *dpp*D3xCi^*KO*^ has been modelled independent of external factors it is expressed at basal level anywhere. Cells expressing *dpp*D3xCi^*ptc*^ more than the basal level are in the NAC range. The expression of both genes in the wing imaginal disk is also indicated by using coloured bars. The blue circle inside the bar, indicate the position of a cell expressing *dpp*D3xCi^*ptc*^ at the basal level. The color scale used in these bars is shown in **C)** black meaning no expression (0*nM/min*), and full color meaning high expression (1*nM/min*). The inset in **B)** depicts the activator/repressors (CiA/CiR) gradients generated by Hh signalling: activator concentrations are higher close to the Anterior/Posterior border.

From the analysis of these mathematical expressions, we can estimate which biochemical characteristics can modify the spatial expression of two genes controlled by the same TFs and in what way. The experimental evidences that motivate our analysis are mainly related with the affinity and the number *n* of the binding sites. By electrophoretic mobility shift assays Parker and coauthors found in [13] that Ci binding sites in the *ptc* enhancer have considerably higher affinity than *dpp* sites. The same authors constructed transgenic fly lines that allow them to compare the transcriptional activity of reporter genes containing different variants of these sites modifying their affinity.

Since there are two opposing signals we have to determine when a cell is net activated or repressed. These notions have been adopted from [13,16] where using reporter genes, the activity of different versions of the *dpp* enhancer containing three low-affinity sites (*dpp*D-Ci^*wt*^), three high-affinity sites (*dpp*D-3xCi^*ptc*^) or three null-affinity sites (*dpp*D-3xCi^*KO*^) were compared. The reporter gene with null-affinity sites provided the basal expression, since it reflects the effects of all other factors which are different than Ci on *dpp*. Specifically, in [13,16] the effects of Ci signalling with low- or high-affinity enhancers was measured comparing the gene activity *versus* the basal in any cell. Cells expressing a gene with higher expression rates than the basal level are called net activated cells. The set of all the net activated cells constitutes the *net activated cellular* (NAC) range. Fig. 1 shows how this range is determined by using a thermodynamic model in the same way as was done from measurements in [13].

We want to find out which concentrations of activators and repressors, [CiA] and [CiR], will provide more or less gene expression than the basal. So, we define a threshold separating concentrations of both TFs that would produce net activated cells or net repressed cells using the BEWARE operators. Then, we can use the threshold between net activation/repression concentrations to determine the limit between the ranges of net activated or repressed cells. Once we have defined the ranges of activated cells we can predict how biochemical differences will affect them as well as the signal intensity. To do this analysis we need to assume that the opposing activator and repressor gradients are monotone and do not change over time (see Fig.1 (B) for a graphical example). Let us mention that the approach we follow is independent of the specific values adopted by the TFs concentrations. Mathematical analysis is able to detect how biochemical differences provoke variations in transcription rates by only assuming that they respond to the same (unknown) opposing activator/repressor concentrations. This is in contrast with the great variability in TFs gradients determination exhibited by the theoretical models fitted in [13]. This methodology is explained in detail in Sec. Methods. Note that another theoretical approach has been proposed in the previous work [23].

With the help of the analysis performed we can now determine which biochemical factors could be involved in the differential expression observed in Hh target genes. To do this we test and compare our theoretical results with the already existent experimental evidence.

### Experiment 1: Transcriptional effects of the reduction of binding sites

In [13], a transgenic fly line carrying a GFP reporter with a single high-affinity Ci site (*dpp*D-1xCi^*ptc*^) was also constructed. In this case, the range of net activated cells was wider than the range of net activated cells for (*dpp*D-3xCi^*ptc*^) and in consequence a broader, but attenuated, expression was observed for the single enhancer case than in the 3 high-affinity sites case. This comparison can be reproduced by using the BEWARE operators. It can be theoretically proved that the cooperativity between the

TFs determines the effect of the binding sites reduction:

- In the presence of total cooperativity or non-cooperativity between the TFs the NAC range would essentially remain unaltered although a reduction of signal intensity could be observed, that is, less repression/activation in the repressed/activated cells.
- If the activators CiA only cooperate between them a reduction in the NAC range would be observed, so some net activated cells for *dpp*D-3xCi^*ptc*^ would change to be net repressed for the gene *dpp*D-1xCi^*ptc*^.
- Finally, in the case of cooperativity only between repressors the NAC range would be incremented, in concordance with the measurements for *dpp*D-3xCi^*ptc*^ and *dpp*D-1xCi^*ptc*^ obtained in [13].

These results (sumarised in Table 1 row **3**)) are balancing a twofold consequence of the reduction in the number of enhancers from 3 to 1. At one hand, the reduction implies the vanishing of any possible kind of cooperativity between TFs. In the case of total cooperativity between TFs these relations are symmetric for activators and repressors, so their disappearance reduces the signalling, that is less transcription in activated cells and more transcription in repressed cells, but it does not modify the balance between net activated or repressed cellular ranges. In the case of asymmetric cooperativity, that is, partial cooperativity either only between activators or repressors, the cooperative specie is loosing that advantage. This would provoke a global reduction of activation, in the case of activators cooperativity, and repression in the case of repressor cooperativity. This forces the NAC range variation: when the cooperativity between activators is removed the NAC range is reduced and it increases when the repressor cooperativity is abolished. The interpretation of these assertions on the contrary allow us to affirm that one of the roles of asymmetric cooperativity between activators/repressors is to increase/decrease the NAC range with respect to the NAC range in the non cooperative case. A summary of these results can be found in Table 1 row **2**). On the other hand, regardless of the cooperativity, the reduction in the number of enhancers also implies that signalling has to be weakened. Both considerations explains the theoretical transcriptional effects of the binding sites reduction.

**Table 1.**
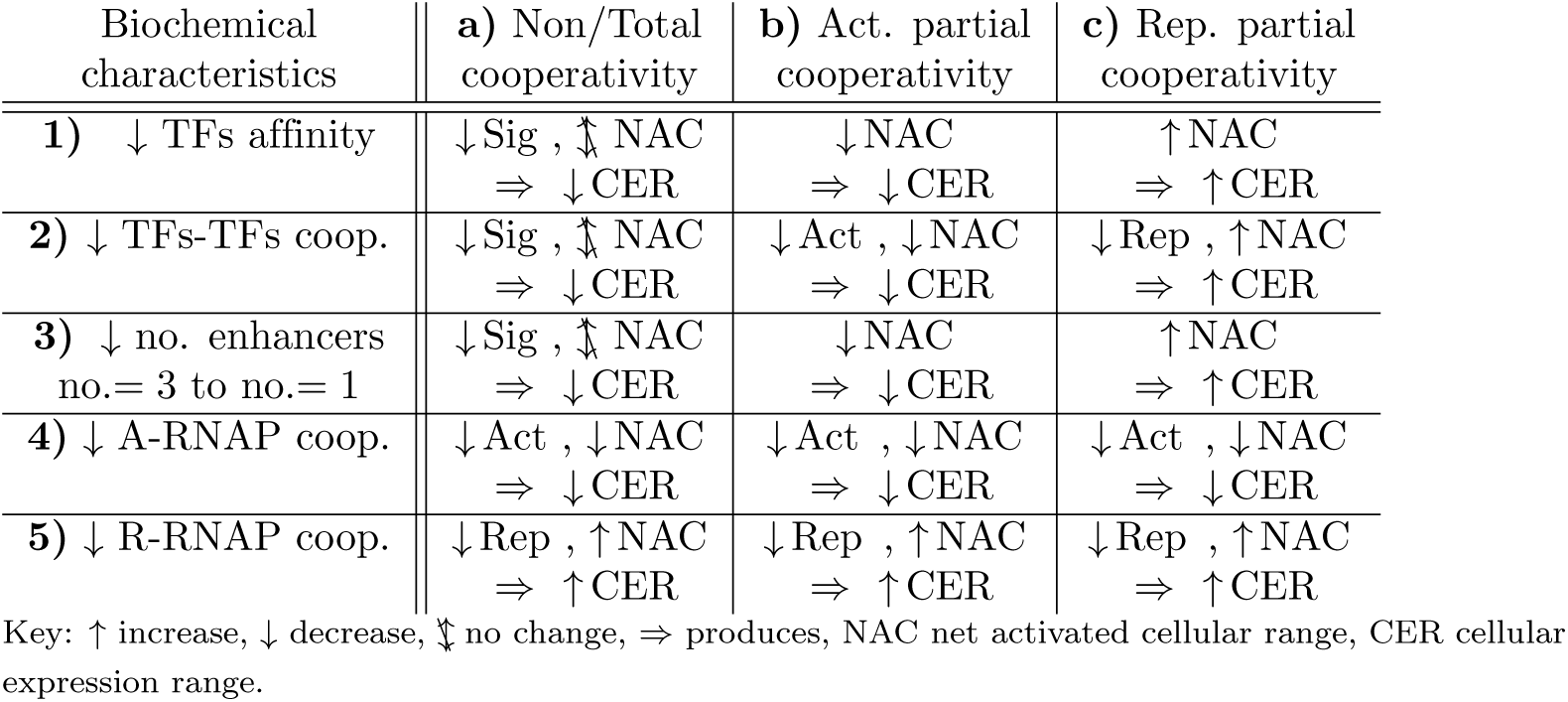
Transcriptional logics in the presence of opposing A/R gradients. This is a simplified comparison of the transcriptional response to different biochemical characteristics between two genes controlled by opposing activator/repressor gradients. The column headings are the kind of TF cooperativity analysed: non/total cooperativity (TFs can cooperate with any other TF), partial cooperativity only between activators or partial cooperativity only between repressors. The biochemical characteristics are: row 1): affinity of TFs for their binding sites, rows 2): cooperativity between TFS, row 3): number of enhancers, row 4) and 5): cooperativity between TFs and RNAP. The variation of affinity considered in row 1) is proportionally equivalent for both activators and repressors. The table shows whether the cellular expression ranges increase or decrease (↑ CER, ↓CER) and how it works.Decreases in cooperativity between TFs and RNAP, rows 4) and 5), again produce globally higher/lower expression rates which cause the increase/decrease in the net activated cellular range (↑ NAC, ↓ NAC resp.) and CER. The response to differences in the other analysed biochemical characteristics varies depending on the kind of cooperation between TFs. If activators and repressor do not cooperate or cooperate globally the net activated cellular region remains unaltered (:↕ NAC) and the signal is weakened (↓ Sig) provoking the decrease of activation in net activated cells but also repression in net repressed cells. On the other hand, if partial cooperation occurs between TFs the same biochemical characteristics can produce either increase or decrease of the net activated cellular region (⇅NAC) and in consequence broader or narrower expression ranges (⇅CER) depending on the type of cooperation.

Fig. 2 provides particular examples of the different behaviours in the NAC range under the same experiment in presence of the cooperativities previously mentioned. Under our modelling, the only transcriptional logic compatible with the experiments is the one which occurs in presence of partial cooperativity between repressors. Nevertheless we can find more concordances in this direction by using other experimental evidences.

**Fig 2.**
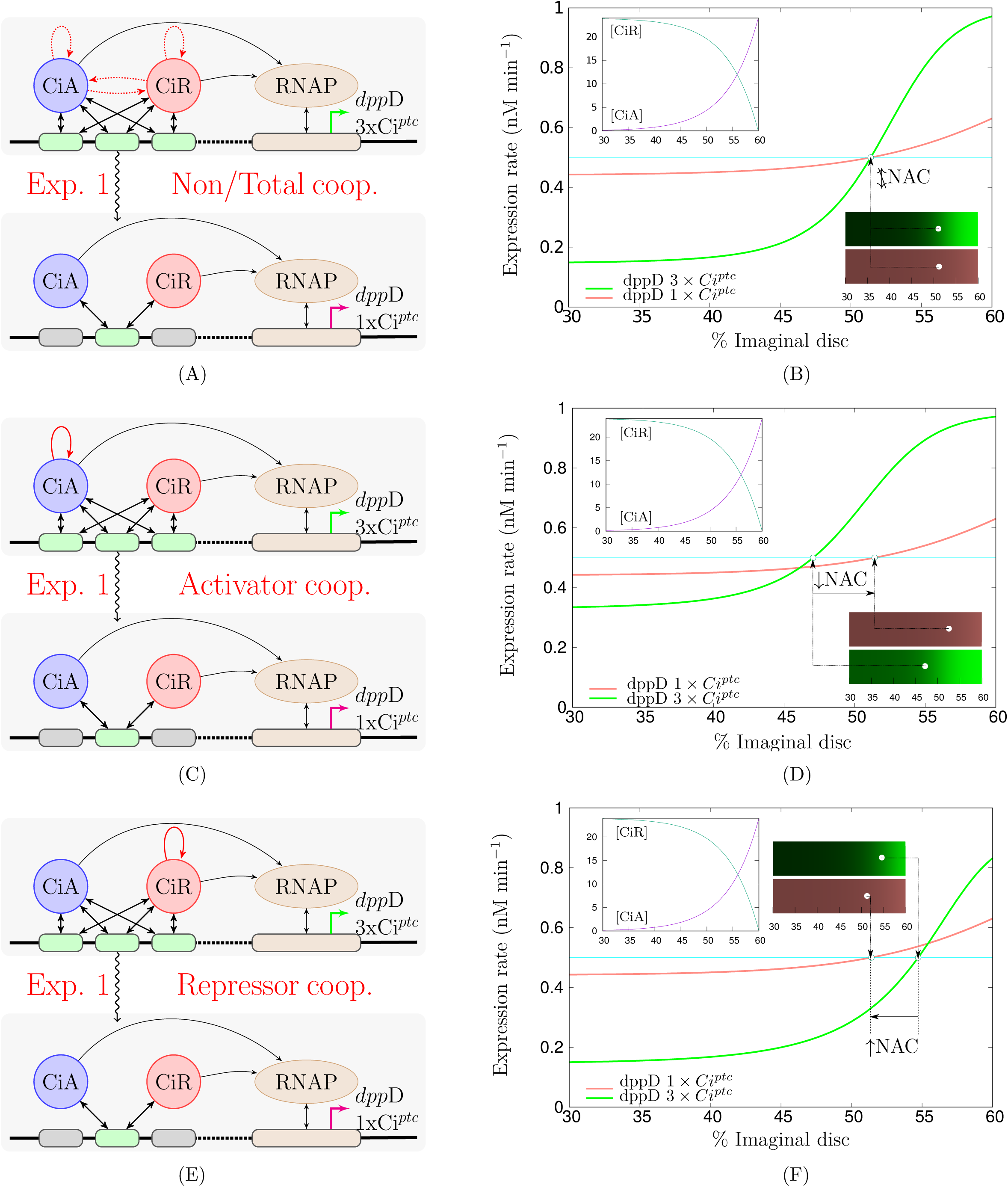
Transcriptional responses to the experiment 1. First column: schematic of the experiment 1: comparison of the expression ranges of reporter genes with 3 high-affinity sites (*dpp*D-3xCi^*ptc*^) and a single high-affinity Ci site (*dpp*D-1xCi^*ptc*^). **A)** corresponds to the non/total cooperativity case where, if cooperativity holds, all the TFs cooperate between them, **C)** to the activators cooperativity case, only activators cooperate, and finally **E)** to the repressors cooperativity case where only repressors cooperate. Second column shows the different transcriptional responses that can be theoretically described depending on the case of cooperativity considered. The schemes and plots employ the same keys explained in Fig. 1.

### Experiment 2: Differential affinity effects

The second experiment we focus on is the comparison of net transcriptional rates of reporter genes containing either three high-affinity sites version of the *ptc* enhancer (*dpp*D-3xCi^*ptc*^) or three low-affinity *dpp* sites (*dpp*D-Ci^*wt*^). It was observed that higher Ci affinity provides a reduction in the net activated cellular region, that is a relevant intermediate region where net activated cells for (*dpp*D-Ci^*wt*^) are net repressed for (*dpp*D-3xCi^*ptc*^). It was also observed that increased affinity provides stronger activation in the region close to the A/P border as well as a stronger repression in regions far from the same border (see Fig.2 D in [13]).

If we accept that the cooperativity between TFs is working in the same way in the binding to both, *dpp* and *ptc*, versions of the binding sites, our analysis suggests that the effect of the reduction in affinity could again depend on the type of cooperativity occurring between TFs:

- In the presence of total cooperativity or non-cooperativity between the TFs the NAC range would essentially remain unaltered although a reduction of signal intensity could be observed, that is, less repression/activation in the net repressed/activated cells.
- If the activators CiA only cooperate between them the NAC range would be reduced, because this range can be proved to be monotone increasing with affinity, that is, the more affinity the wider the NAC range.
- In the case of cooperativity only between repressors the NAC range would be incremented, because this range is monotone decreasing with affinity, that is, the more the affinity the narrower the NAC range.

Again the repressor cooperative model is the only in concordance with the results observed for *dpp*D-3xCi^*ptc*^ and *dpp*D-3xCi^*wt*^ in [13]. These theoretical behaviours can be seen in Fig. 3. In all the previous analysis, summarised in Table 1 row 1), it was considered that the change in affinity of the enhancers affect to both, activators and repressors, in a proportional manner. The results for partial cooperativity also requires that the addition of the concentrations of activators and repressors at any cell is the same for any cell of the Anterior compartment (total amount of Ci conservation). See Sec. Results for details.

**Fig 3.**
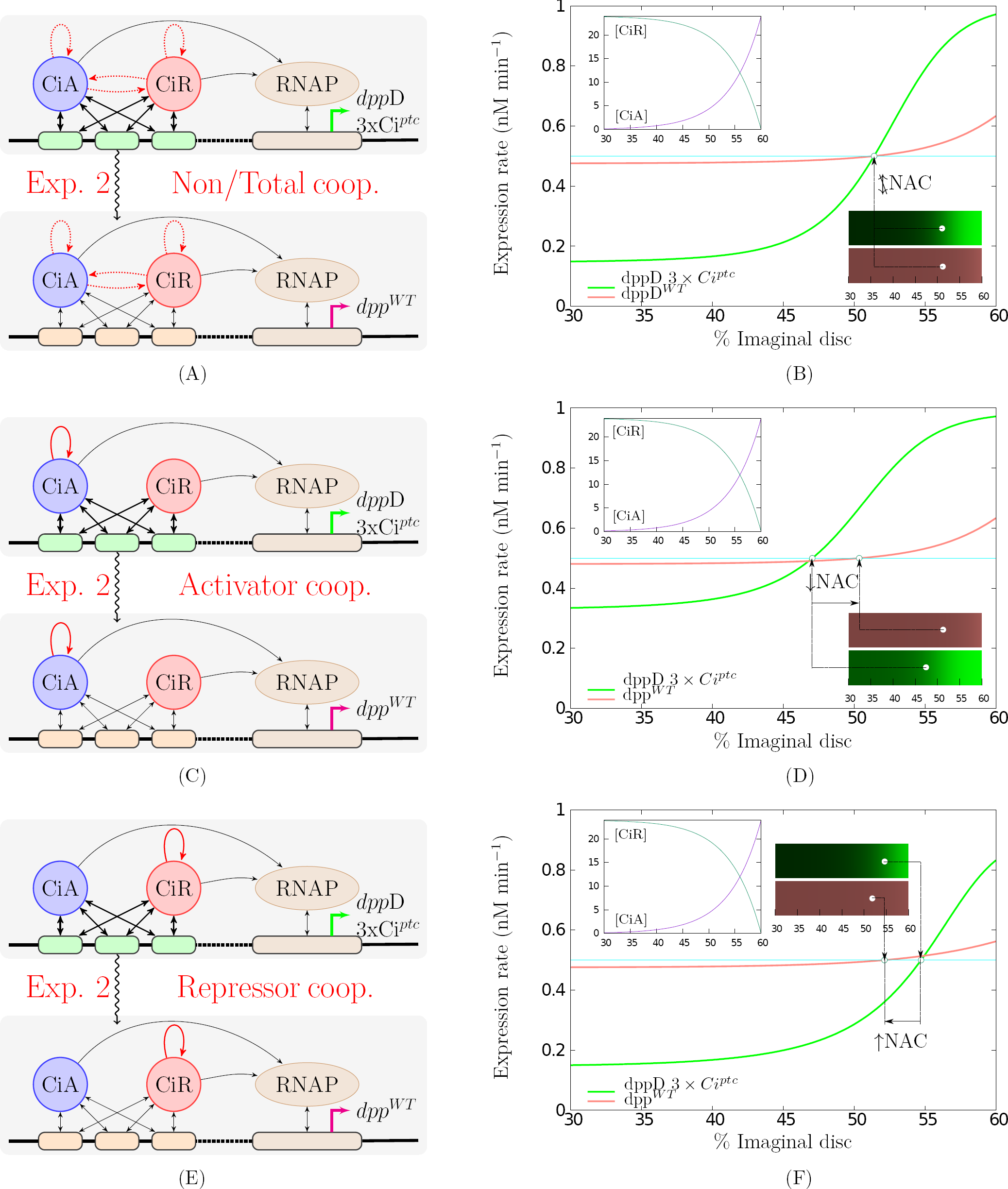
Transcriptional responses to the experiment 2. Figures in the first column: schematic of the experiment 2: comparison of the expression ranges of reporter genes with 3 high-affinity sites *dpp*D-3xCi^*ptc*^ or 3 low-affinity site *dpp*D-3xCi^*wt*^. TF_s_-DNA binding site affinities are indicated by thicker or thinner double-headed straight arrows. **A)** correspond to the non or total cooperativity case where, if cooperativity holds, all the TFs cooperate between them, **C)** to the activators cooperativity case, only activators cooperate, and finally **E)** to the repressors cooperativity case where only repressors cooperate. Figures in the second column shows the transcriptional responses that can be theoretically described depending on the case of cooperativity considered. The schemes and plots employ the same keys explained in Fig. 1

Thus, in the case of the Hh target genes, both results are only compatible with the presence of cooperation between repressors. This conclusion coincides with the one obtained in [13] by using numerical fittings to experimental data and with [16] where the authors found that *dpp* requires low-affinity binding sites for normal activation in regions of low Hh signalling. Indeed, our analysis allow us to interpret the roles of differential affinity and repressor cooperativity for the Hh target genes in the following terms:

- The analysis of experiment 1 show us that repressors cooperativity reduces the NAC ranges of both genes with respect to the non-cooperativity case.
- Furthermore, the analysis of experiment 2 implies that this reduction of the NAC region is less effective with low affinity binding sites of *dpp* than with high affinity binding sites of *ptc*.

Moreover, the previous point of view would imply that the NAC range for *dpp*D-1xCi^*ptc*^ should contain to the NAC range for *dpp*D-3xCi^*wt*^, and the latest should contain the NAC range for *dpp*D-3xCi^*ptc*^. In fact, this relations are fully compatible with the results of [13]. More concretely, in Figures 2 and 4 [13] it can be observed that the NAC ranges for *dpp*D-1xCi^*ptc*^, *dpp*D-Ciwt and *dpp*D-3xCi^*ptc*^ occupy from the 43%, 49% and 54% of the disc width respectively to the A/P border (which is around to the 60% of the disc width).

### Transcription logics in the presence of opposing transcription factors

Although the results of our analysis have been before directly applied to the Hh target genes, the analysis can be performed for any other genes controlled by opposing transcription factors, A and R, activators and repressors respectively. We will adopt from now on this general approach, and in the particular case of Hh target genes CiA and CiR will play the role of A and R. As it was pointed out in the Introduction the previous results are not compatible with the transcriptional logic of the activator threshold model. So, in this section we will describe the versions of the transcriptional logic that could be found depending on the cooperativity between the TFs: non/total cooperativity, activators partial cooperativity and repressors partial cooperativity.

The results of our analysis are summarised in Table 1. It shows the relative size of the cellular expression ranges (CERs) of two genes, *gene1* and *gene2*, controlled by the same opposite TF gradients although exhibiting some differences in their biochemical characteristics. The considered biochemical differences between *gene1* and *gene2* are listed in rows:

1. ↓ TFs affinity: the affinity for the binding sites of *gene1* is smaller than the affinity of the same TFs for the binding sites of *gene2*. Following [13] this occurs, for instance, in the case of *dpp*D-3xCi^*wt*^ and *dpp*D-3xCi^*ptc*^. The affinity decrease considered is proportional for A and R,
2. ↓ TFs-TFs coop.: the TFs cooperate in a less intense manner in the binding to the enhancers of *gene1* than in the binding to the enhancers of *gene2*, but both genes have the same type of cooperativity,
3. ↓ no. enhancers: *gene1* has less binding sites than *gene2*, as for instance the genes *dpp*D-1xCi^*ptc*^ and *dpp*D-3xCi^*ptc*^,
4. ↓ A-RNAP coop.: the activator A is a weaker transcriptional activator for *gene1* than for *gene2*, exhibiting weaker cooperativity between the activators and the RNA polymerase,
5. ↓ R-RNAP coop.: the repressor R is a weaker transcriptional repressor for *gene1* than for *gene2*, exhibiting weaker anti-cooperativity between the repressors and the RNA polymerase.

The results compiled in Table 1 also summarise the way in which the CER variations happen. ↑ CER / ↓CER indicate *gene1* has a broader/narrower CER than *gene2* respectively. These CER variations depends on signal modulation as well as on the NAC range variations. In this sense the theoretical model suggest that the following situations can occur:

- ↓ Act: Global decrement of the expression rates of *gene1* with respect of those of *gene2*,
- ↓ Rep: Global increment of the expression rates of *gene1* with respect of those of *gene2*,
- ↓ Sig: Signalling decrement, that is, for *gene1* lower expression rates in net activated cells and higher expression rates in net repressed cells.
- ↓ NAC: decrement of the NAC range, when the NAC range for *gene1* is smaller than the NAC range for *gene2*,
- ↑ NAC: increment of the net activated cellular range of *gene1* with respect to the NAC range of *gene2*,
- 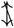 NAC: when the NAC ranges for *gene1* and *gene2* are equal.

The consequences of the differences in biochemical characteristics listed in Table 1 clearly justify the existence of several versions of the transcriptional logic in the presence of opposing gradients depending on the kind of cooperativity between TFs. The region of net activated cells is highly relevant in the Non/Total cooperativity case because the effects of some changes in biochemical characteristics only weaken the signalling (↓Sig) and in consequence they do not modify the NAC range 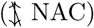 (see Fig. 4(B), (C), (D) in Supplementary Material). In the case of partial cooperativity only between repressors or activators global increment or decrement of expression rates can be deduced which clearly provoke increase or decrease of net activated cellular region (see Fig. 6(D), Fig. 7(D) in Suplemental Material). However, not always a variation in NAC range is due to a global increment or decrement of expression rates as can be seen in Fig. 5(B) and 6(B). Nevertheless, we can observe that the consequences resulting from some others biochemical characteristics, as those described in Table 1 rows 4), 5), are qualitatively the same, independently of cooperativity. Table 1 has been introduced for the sake of clarity, although it does not cover all the relationships and properties deduced in our analysis. Figures 4, 5 and 6 in Supplementary Material illustrate the results stated in Table 1 rows 1), 2), 3). In these graphs thresholds and transcription rates are represented in magenta for *gene1* and green for *gene2* in order to appreciate relative differences.

The same BEWARE operators can be used to find out the expression rates for a single activator gradient simply by setting to zero the concentrations of repressors. In the same way, we can also perform the analysis of the same kind of experiments and the results are in concordance with transcriptional logic of the activator threshold model. See Supplementary Material 1.4.

## Discussion

In this work we attempt to deepen the current understanding of genetic expression. By using a theoretical approach we are able to isolate the biochemical mechanisms involved in the expression of genes which operate through opposing activator and repressor TF gradients, and the degree of their involvement. The BEWARE method is a well accepted modelling tool which allows us to represent the delicate balance between opposing signals using mathematical expressions and see how these proportions are affected by the other biochemical characteristics involved. By reducing the previously long BEWARE formulae into compact mathematical expressions we are able to deduce the existence of several different forms of transcription logic, that is, several scenarios where the same biochemical characteristics between genes produce absolutely different consequences at a tissular level. The detailed description of the different scenarios and the relationship between them allows us to contrast this theoretical framework with the experimental evidence obtained in specific systems. This has been achieved in the case of the Hedgehog Target genes *dpp* and *ptc* where we obtain conclusions analogous to those obtained in previous work using other techniques.

This work extends the applicability of the BEWARE method since the relevant qualitative information can be extracted from the compact models. The fact that these models could be applied in a similar way to other biological systems means there are many interesting implications beyond the scope of this paper.

## Methods

### Deduction of the BEWARE operator for two opposing TF gradients

As a first step, we apply the ideas of the statistical thermodynamic method to a gene, *g*, controlled by two opposing transcription factors {*A, R*}, activator and repressor respectively. Our goal here is to deduce expressions for the change of the concentration of protein G over time in terms of the concentrations of A and R, [*A*] and [*R*], i.e.,

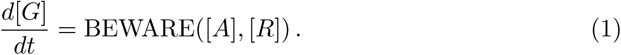

Here ‘BEWARE()’ represents a mathematical function specifying the dependence with respect to the activation/repression role of the TFs. This is independent of other possible factors relevant for the protein evolution as for instance degradation or spatial dispersion. In the model, the binding reactions of TFs and RNAP in the enhancers and promoter, respectively, are much faster than the synthesis of the protein G, hence it will be considered in thermodynamic equilibrium given by the Law of Mass Action. If *B* is an empty regulatory region, a set of non occupied enhancers-promoter, the complexes *BA, BR* and *BRNAP* have concentration at equilibrium given by

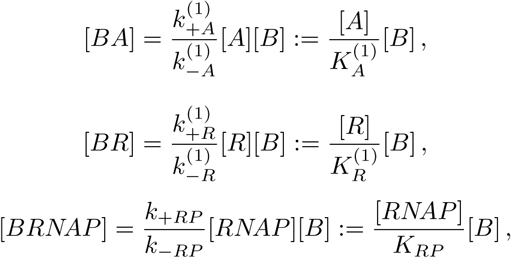

where 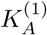, 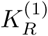, *K*_*RP*_, [*A*], [*R*] and [*RNAP*] are dissociation constants and concentrations of activators, repressors and RNA polymerase. So the quotients, 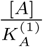, 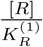, and 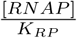, are dimensionless. The superscript (1) stands for the dissociation constant of a reaction that takes place in absence of another TF, previously bound to another enhancer (note that, since the sets only have one promoter, the superscript is not needed for the RNAP dissociation constant). Let us observe that the higher the affinity between a protein/complex and the binding site the lower the dissociation constant in the corresponding binding reaction. The consecutive binding of more that one transcription factor is considered as a sequential and competitive process, such that the reactions

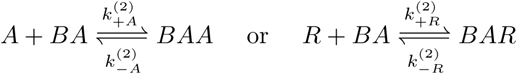

are given by equilibrium concentrations

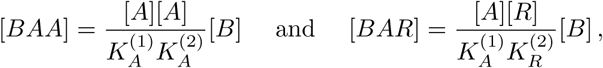

here, the superscript (2) denotes the dissociation constant for a reaction of a TF that already binds the operator with one TF in another site. The competition is modelled such that the dissociation constant of the free sites configuration does not depend on their position, but might depend on other TFs already bound to the same set of enhancers by cooperativity or anti-cooperativity.

We denote as non cooperative TF_s_, all those proteins whose enhancer affinity is not modified by any previously bound TF_s,_ that is, they verify 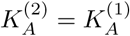 and 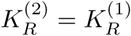. This assumption implies sequential independence of the equilibrium concentrations since [*BRA*] = [*BAR*]. It is plausible to assume the same relation for later bindings, that is, 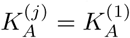 and 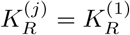 for *j* ≥ 2. In consequence of this sequential independence we denote the dissociation constants as *K* _*A*_ and K_*R*_ omitting the superscript. So, if all the TFs under consideration are non cooperative we easily deduce that the concentration at equilibrium, of a configuration with *j*_*A*_ activators and *j*_*R*_ repressors bound, is

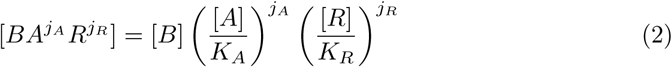

independently of the sequential order of binding and of the specific positions occupied by the TFs. Although *Drosophila*’s wild type cis-regulatory elements involve a total number of 3 binding sites we are going to compare with experiments where these binding sites have been reduced to 1. Thus, we will consider in our model n ≥ 1 the number of TFs binding sites. In all cases, we have a restriction for the possible number of bound transcription factors. So, *j*_*A*_ + *j*_*R*_ ≤ *n* has to be verified, and in consequence *j*_0_ = *n* – *j*_*A*_ – *j*_*R*_ ≥ 0 denotes the number of free spaces in the configuration.

On the other hand, cooperativity occurs when the existence of other previously bound proteins affects the affinity of the new binding protein of type *i, k* = *A, R*, that is:

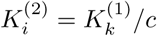

where *c* is a positive constant greater than 1 if proteins cooperate, and less than 1 if anti-cooperativity occurs. Since the only difference between cooperativity and anti-cooperativity is a threshold value for c, in the subsequent modelling we will refer to the constant *c* and not distinguish between both cases. If cooperativity occurs it would be necessary to know which TFs are affected by other TFs since the equilibrium concentration will depend on these relationships. In previous literature, total and partial cooperativity have recently been proposed to play an important role in the Hh/Shh target genes by means of the Ci/Gli TFs [10,13,23]. Partial cooperativity of the activators would occur when the existence of a bound activator modifies equally the affinity of any posterior activator binding, that is 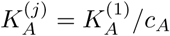 for *j* ≥ 2. The same applies for repressors. Total cooperativity would occur when the presence of a bound TF modifies the affinity of any posterior binding in the same manner, i.e. 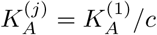 and simultaneously 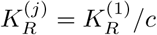 for *j* ≥ 2 (see for instance [11]). From now on, we will denote the activator and repressor dissociation constants as *K* _*A*_ or *K*_*R*_, such that

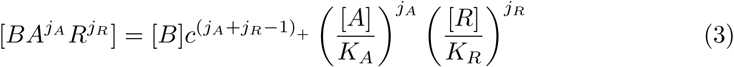

in the presence of total cooperativity, while

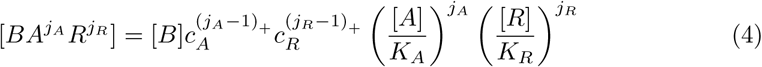

if partial cooperativity for TFs occurs. Here, (·)_+_ denotes the positive part function ((*x*)_+_ = *x* if *x* > 0 and (*x*)_+_ = 0 if *x* ≤ 0). This is needed because the cooperativity will not take place unless two or more cooperative TFs are present in the configuration. In the rest of this paper, we will designate the cases when the TFs cooperate between them totally and partially as {{*A*, *R*}_*c*_} and {{*A*}_*CA*_, {*R*}_*CR*_ } respectively. Note that this notation covers the case of non cooperativity since it would correspond to the case {{*A*, *R*}1} or equivalently {{*A*}_1_, {*R*}_1_}.

The binding sites are ordered spatially and, in general, there is not a unique spatial distribution for a configuration with *j*_*A*_ activators, *j*_*R*_ repressors and *n – j*_*A*_ *– j*_*R*_ free sites. For instance, if we consider *j*_*A*_ = *j*_*R*_ = 1 there are six possible spatial distributions with the same elements *(ARO, RAO, AOR, ROA, OAR, ORA* where *O* denotes the empty space). In our description, spatial localisation of bound particles is not considered. In fact, for a specific configuration with *J*_*A*_ activators, *j*_*R*_ repressors and *j*_*0*_ free sites 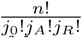 different spatial configurations are plausible, where k! denotes the factorial of *k*.

Regarding the promoter’s RNA polymerase binding process, the TFs work together trying to promote or repress the binding process [12] by a mechanism known as recruitment [14,15]. Thus, we consider that the activators interact with RNAP with ‘adhesive’ interaction [4] that gives rise to a modification of the RNA polymerase binding affinity: *K*_*RP*_*/a*^*j*_*A*_^ where *a* is a cooperativity constant greater than 1. In contrast, the effect of *j*_*R*_ repressors is modelled in terms of a ‘repulsive’ interaction that modifies the binding affinity *K*_*RP*_*/r*^*j*_*R*_^ with an anti-cooperativity factor *r* < 1 (repressor interaction). We will refer to these parameters as TF transcriptional activation/repression intensity.

By using the previous guidelines we will now describe the concentrations of all possible configurations as was done in [2,19]:

### Step 1: Construction of the sample space

All the possible ways of obtaining an equilibrium concentration with *j*_*A*_, *j*_*R*_ and *j*_*P*_ activators, repressors and RNA polymerases is given by the states

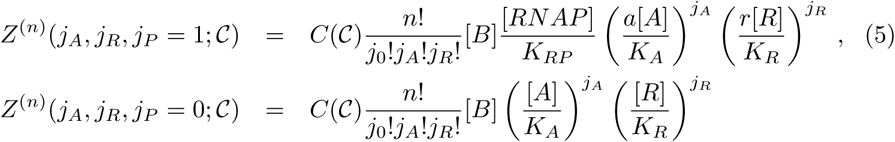

where *j*_*P*_ = 1 means there is a bound RNA polymerase and *j*_*P*_ =0 there is none, *j*_*0*_ = *n – j*_*A*_ *– j*_*R*_ ≥ 0, and the variable C describes the relation of cooperativity between the TFs. Specifically, by using (3) and (4), the cooperativity function *C* takes the values

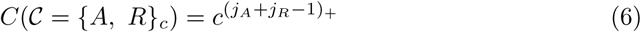

and

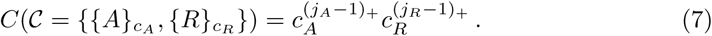

This allows us to describe the entire sample space, i.e. the space of all the possible configurations, by

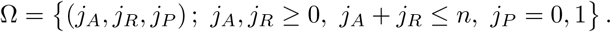

### Step 2: Definition of the probability

Once we have described all the possible configurations in terms of the concentrations of activator, repressor and RNA polymerase, we easily obtain the probability of finding the promoter in a particular configuration of *j*_*P*_ RNA polymerase and *j*_*A*_, *j*_*R*_ TFs related by a cooperativity relation C as

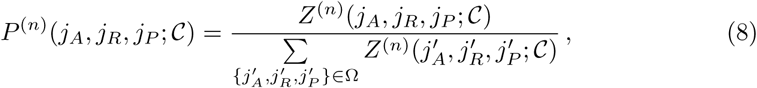

for all *(j*_*A*_, *j*_*R*_, *j*_*p*_) ϵ Ω.

### Step 3: Definition of the BEWARE operator

In this last step, the BEWARE operator is obtained in terms of the probabilities *P*^(*n*)^. Following the work of Shea et al [19] the synthesis of a certain protein depends on the total probability of finding RNA polymerase in the promoter, specifically, the synthesis is proportional to the marginal distribution of the case *j*_*P*_ = 1 [4,7,13]. This justifies the definition of the BEWARE operator as

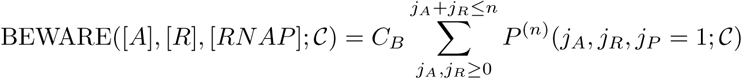

where in definition (8) expression (5) is assumed and *C*_*B*_ is a proportionality constant that could depend on other factors not considered in this work. Splitting the denominator in two sums, when RNA polymerase is bound or not bound to the configuration, this expression can be rewritten in terms of the regulation factor function, *F*_*reg*_:

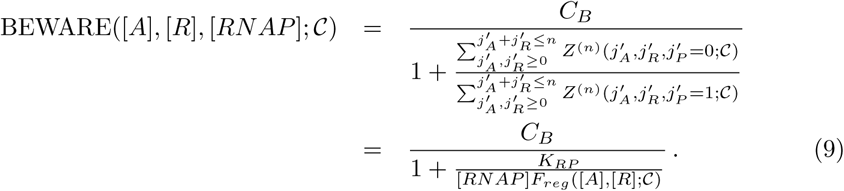

Doing some basic algebra, this regulation factor can be reduced to facilitate the understanding of the general process (see Supplementary Material 1.1). This has been done by using a classic strategy employed for obtaining the General Binding Equation more than a century ago [5]. This, have not yet been applied, in this context, to the authors knowledge. In fact, we can prove that the regulation factor can be equivalently written as

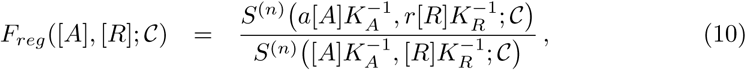

where the explicit expression of *S*^(*n*)^ (*x*, *y*; *C)* depends on the kind of cooperativity presumed, that is

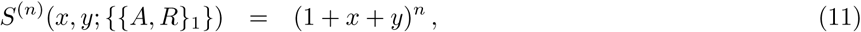

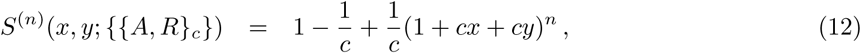

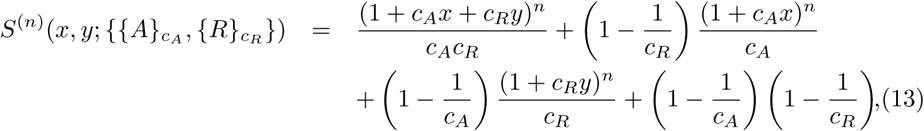

for the non cooperative, total and partial cooperative cases respectively. Note that the mathematical complexity in these expressions is mainly related to the assumed cooperativity.

### Versions of transcription logic in the presence of opposing gradients

In this section we are going to describe what the transcriptional reaction of genes, controlled by the same opposing TFs, would be when there are biochemical differences between them. As we explained in the section Results the consequences of such differences will depend on the type of cooperativity occurring between the TFs. The analysis of the case of single gradients can be found in Supplementary Material 1.4.

### Transcriptional logic in the case of opposing gradients and non/total cooperativity between TFs

We observe that expression (12) coincides with (11) when *c* = 1 which allows us to use the same mathematical expression for both cases, non cooperativity and total cooperativity. So, in both cases the transcription rates are given by (9) with

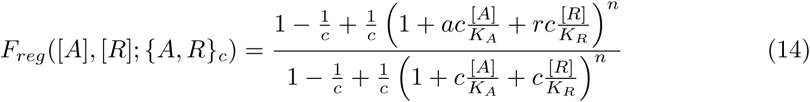

being the regulation factor, where *c* = 1 if there is no cooperativity between TFs and *c >* 1 total cooperativity occurs. In fact, we prove that the transcription logic will be basically the same in both cases.

### Determination of net activation/repression concentrations and cell activated ranges

Thanks to the BEWARE operator we can theoretically describe which concentrations of activators and repressors will cause higher or lower gene expressions than the basal level, that is, we can describe in great detail the effect of the balance of both signals. Note that the basal state, determined by the absence of TFs, that is [*A*] *=* [*R*] = 0, corresponds to *F*_*reg*_ = 1 in expression (9). Thus, the regulation factor describes an effective increase (for *F*_*reg*_ > 1) or decrease (for *F*_*reg*_ *<* 1) of the number of RNAP molecules bound to the promoter, with respect to the basal level, as was stablished in [4].

We can see that in the case of (14) the threshold between activation/repression concentrations (that is *F*_*reg*_ = 1) is determined by the linear relation

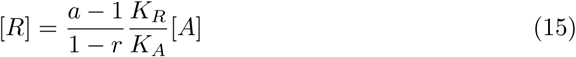

dividing the plane [*A*] – [*R*] into two parts that we can denominate activation region if 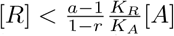 and repression region if on the contrary 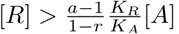. See Fig. 4 (A) in Suplemental Material where examples of these thresholds are depicted for different values of the parameters. Obviously, the threshold (15) is a linear relation between concentrations of activators and repressors. The steepness of this straight line is: independent of the values *c* and *n*, depends on the TFs affinities through the ratio *K*_*R*_*/K*_*A*_, it increases with respect to *a* and decreases with respect to *r* (since *r* < 1). This justifies the behaviour of the net activated cellular ranges in the case of non/total cooperativity stated in Table 1 as we will now explain.

We can take this information and by considering appropriate gradients of activators and repressors we can define the tissular regions of net activated and net repressed cells, which are, regions of cells expressing more or less than the basal expression level (see Fig. 1 for a detailed explanation). For the sake of clarity, and taking into account that our main goal is to understand how these mechanisms could modify the expression of the Hh target genes, we are going to assume that transcription factors act in the same way as Cubitus works in the *Drosophila* system. Hh secreted from the posterior into the anterior compartment of the wing imaginal disc results in opposing gradients of activator and repressor Ci. The A/P boundary is located at around 60% of the dorso-ventral (D/V) axis. The influence of Hh gradient can be appreciated in the middle of the anterior compartment, specifically the cells located in the region between the 30% and 60% on the D/V axis, approximately. In order to model these TFs concentration distributions we are going to assume that they do not change over time and both are monotonic along the tissue,

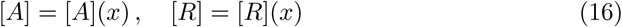

being [*A*](*x*) decreasing and [*R*](*x*) increasing in terms of *x*, the distance from the A/P boundary. See example in inset in Fig. 1 (B) where the [CiA]/[CiR] decreases/increases from the A/P border. The subtle balance between opposing signals involved in the variation of the binding sites affinities and its consequences requires of an extra hypothesis on the TFs gradients: the conservation of the total amount of TFs, i.e.

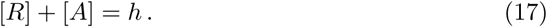

Hence the concentrations will be restricted to a straight line in the [*A*] – [*R*] plane (see Fig. 4 (A), and Fig. 5 and 6 (A), (C), (E)). Insets in Figs. 1, 2, 3 and 4, 5 of Supplementary Material, show the distributions [*A*], [*R*] proposed in [13] corresponding to

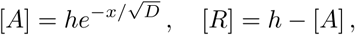

being *x* the distance from the A/P boundary, *h* the TFs total concentration and *D* is the steepness of the gradient. The intersection points between the straight line (17) and the thresholds, ([*A*]_*th*_, [*R*]*_th_)* (represented by black circles in Suplemental Material Figs. 4 (A), and Fig. 5 and 6 (A), (C), (E)) will determine a boundary between genetically activated and repressed cells. That is, repressed cells will be those containing concentrations ([*A*], [*R*]), verifying (17) and [*A*] < [*A*]_*th*_. For activated cells this would be [*A*] > [*A*]_*th*_. In consequence, they would express transcription rates lower/higher than the basal. Due to the monotonic nature of the TFs distributions, (16), activated cells are closer to the A/P boundary and the limit of the percentage of the wing imaginal disc occupied by activated cells will be determined by the distance *x*_*th*_ given by

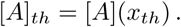

This limit is represented by blue circles in Fig. 4 (B), (C), (D), and Fig. 5 and 6 (B), (D), (F).

### Transcriptional consequences of differential biochemical characteristics

Now, by using previous considerations, we want to justify the transcription logic in presence of total cooperativity or in absence of any cooperativity, results collected in Table 1, column a) and represented in Suplemental Material Fig. 4. In this case it is quite easy to see the behaviour of the net activated cellular range. Equation (15), which determines the threshold between activation/repression concentrations, does not depend on the number of enhancers *n* or the cooperativity *c* and depends on the TFs affinities in terms of the ratio *K*_*R*_*/K*_*A*_. Thus, these regions do not change 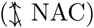 for genes *gene1* and *gene2* such that:

1. The affinity for the binding sites of *gene1* is smaller than the affinity for the binding sites of *gene2* in a proportional manner. In terms of the dissociation constants, this would be expressed as 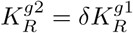 and 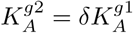 being 0 < *δ* < 1 and this occurs because we are considering proportional change of affinity for activator and repressors, 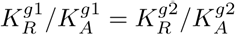.
2. The TFs cooperate in less intense manner for *gene1* than for *gene2*, that is 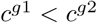.
3. *gene1* has less binding sites than *gene2*, that is 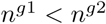.

Here, the superscripts *g*1 and *g*2 stand for the parameters of the genes *gene1* and *gene2*, respectively. However, in the case of differential affinities, where the proportionality is not verified, the net activated cellular range would change. For instance, if 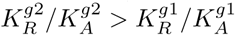 then the net activated cellular range for *gene1* would be narrower than for *gene2*.

The rest of the assertions in Table 1, column a) requiere some simple monotonicity properties that have been checked with Lemma 1.1 and Lemma 1.2 in Supplementary Material. The biochemical differences, numbered **1**), **2**) and **3**), have been proven to verify that

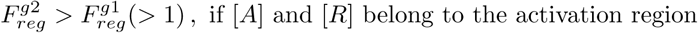

and

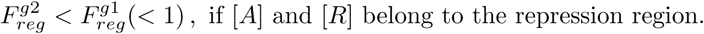

Note that the BEWARE operator is monotonic increasing, with respect to the regulation factor *F_reg_* which allow us to extrapolate these estimates to expression rates. These three results have been interpreted as a signal weakening. Biochemical differences **1**), **2**) and **3**) can not modify the character of net activation/repression but are able to make signalling less efficient. That is, they do not change the NAC range but they cause less activation in the activated region and less repression in the repressed region (↓ Sig). In consequence, we can say that in situations **1**), **2**) and **3**) the expression rates will decrease in the net activated cellular range and increase in the repressed cells, which will attenuate the cellular expression range. See Fig. 4 in Supplementary Material where these variations have been depicted.

In contrast, assertion *3)* in Lemma 1.2 implies that

- If *A* is a weaker transcriptional activator in *gene1* than in *gene2*, exhibiting lower cooperativity between the activators and the RNA polymerase 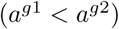, then expression of *gene1* will be smaller than in *gene2* because

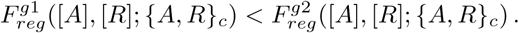 Obviously, this involves globally lower transcription rates (↓ Act) and more restricted net activated cellular ranges (↓ NAC) for *gene1* than for *gene2*.
- If *R* is a weaker transcriptional repressor in *gene1* than in *gene2*, exhibiting lower anti-cooperativity between the repressors and the RNA polymerase 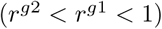, then expression rate of *gene1* will be higher than *gene2* expression because

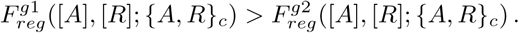 Obviously, this involves globally higher transcription rates (↓ Rep) and wider net activated cellular ranges (↑ NAC) for *gene1* than for *gene2*.

### Transcription logic in the case of partial cooperativity between TFs

In the case of partial cooperativity, the expression rates are given by (9) where the regulation factor is defined by:

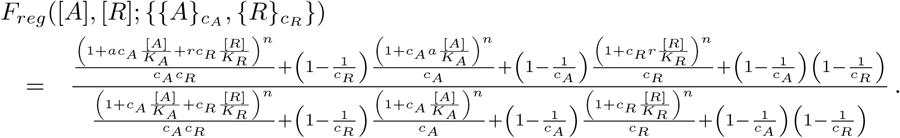

Compared to (14), in this complex representation of the expression rates the activation threshold is not as clear. So we need to do a little more delicate mathematical analysis. In fact, if we impose the threshold equation

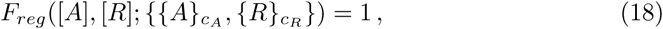

it can be shown that, if *n* = 3 and *c*_*A*_, *c*_*R*_ ≥ 1, this threshold is determined by an unique increasing function *f*, verifying

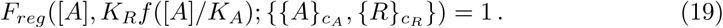

It separates the concentrations [A]-[R] into those which provoke net activation, when [*R*]/*K*_*R*_ < *f* ([*A*]/*K*_*A*_), and those which provoke net repression, when [*R*]/*K*_*R*_ > *f* ([A]/K_*A*_)) (see Supplementary Material 1.3.1 for definition and analysis of the function f). Note that the threshold of a BEWARE operator with partial cooperativity is not, in general, a straight line although it shows a linear asymptotic behaviour for large concentrations (see Fig. 5 and 6 (A), (C) and (E) in Supplementary Material). Note also that the same analysis implies that the function f is independent of the TF affinities: *K*_*A*_ and *K*_*R*_.

We can understand the effects of partial cooperativity clearly in certain limit regimes such as:

- Cooperativity only between repressors, that is *c*_*A*_ = 1 and *c*_*R*_ > 1. The expression rates have been proved to be monotonic decreasing with respect to repressors cooperativity, that is, the more cooperativity the less expression because cooperativity increases repression effectivity (see Lemma 1.3 *2*)). Then, obviously, less cooperativity implies less repression provoking wider NAC and CER.
- Cooperativity only between activators, that is *c*_*A*_ > 1 and *c*_*R*_ = 1. The counterpart results in this other case shows that expression rates are increasing with cooperativity between activators since it increases activation effectivity (see Lemma 1.3 *1*)).Then a reduction in cooperativity between activators reduces NAC and CER.

These result have been summarised in Table1 cells 2b) and 2c) and represented in Fig. 5 (D), Fig. 6 (D).

When both TFs cooperate simultaneously we have compared the thresholds given by functions *f* with the linear relation (15), obtaining a criteria comparison in terms of the variables that follow

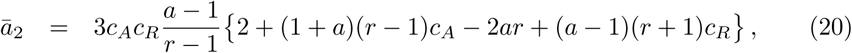

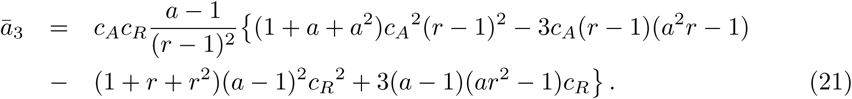

Depending on the positive or negative sign of these values it can be proven that the change from partial cooperativity to non cooperativity causes:

- If 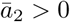, 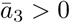: we see an decrement in the net activated cellular range with respect to the non cooperative case.
- If 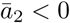, 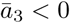,: we see a increment in the net activated cellular range with respect to the non cooperative case.
- In all other cases: An increase or decrease of the net activated cellular range can occur depending on the total amount of TFs considered (*h*), the binding affinities, *K*_*A*_, *K*_*R*_ as well as the activation/repression intensities. A detailed explanation can be found in Supplementary Material 1.3.1.

Another interesting aspect is how these thresholds depend on the affinities of the TFs in presence of partial cooperativity. Let us consider again *gene2* with high affinity binding sites and *gene1* with proportionally low affinity binding sites, that is,

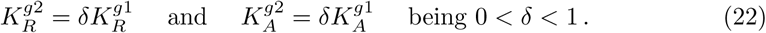

Let us now consider the function *f*. It defines the threshold of a BEWARE operator with partial cooperativity, determined by the cooperativity constants *a, r, c*_*A*_ and *c*_*R*_ and expression (19). This function and the corresponding affinities determine the activator concentration thresholds for *gene1* and *gene2*, 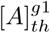 and 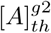, using these expressions

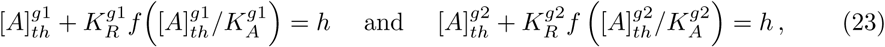

where the conservation of TF concentration, (17), has been considered. Then, under hypothesis (22), the order of 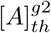 and 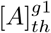 can be determined. It can be proven that when only activators cooperate between them these limit values verify

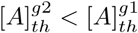

implying that *gene2* has a wider expression range than *gene1*. On the other hand, when only repressors cooperate the inverse relation its true

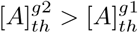

implying that *gene2* has a narrower expression range than *gene1* (see Supplementary Material 1.3.1, Lemma 1.7 for details). In Fig. 3 (D) and (F) both these situations have been depicted when partial cooperativity is either for activators or for repressors. More detailed representations about the thresholds and these range determination process can be found in Figs. 5 and 6 (A) and (B).

Previous results can be generalised to the case of both species cooperating simultaneously and monotonicity has be related with conditions of concavity and convexity of the threshold function *f*. This condition can be understood in terms of the Greater-Than-Additive and Less-Than-Additive effects described in transcriptional activation [7]. The concavity of *f* is the same as observing Greater-Than-Additive effects at the basal level. That is, when *f* is concave, any convex combinations of two different pairs of concentrations, [*A*], [*R*], that give basal expression, will produce net activated cells. On the other hand, when *f* is convex Less-Than-Additive effects can be observed. This result, and our interpretation, are in contrast with the analysis in [23] for the case of cooperative repressors.

The assertions in Table1, cells **4b**) and **5c**) can be verified easily from statement *3)* Lemma 1.3, in Supplementary Material.

## Acknowledgements

O. S. would like to thanks professor J. Garcia-Ojalvo for pointing out reference [4] and R. Bielawski for language editing and proofreading. This work has been partially supported by the MINECO-Feder (Spain) research grant number MTM2014-53406-R, the Junta de Andalucía (Spain) Project FQM 954, and the MINECO (Spain) research grant FPI2015/074837 (M.C.).

## References

1. Aguilar-Hidalgo D, Dominguez-Cejudo MA, Amore G, Brockmann A, Lemos MC, Cordoba A, Casares F. A Hh-driven gene network controls specification, pattern and size of the Drosophila simple eyes. Development 2013;140: 82–92.

2. Ackers GK, Johnson AD, Shea MA. Quantitative model for gene regulation by λ phage repressor. Proc. Natl. Acad. Sci. USA 1982;79: 1129–1133.

3. Ay A, Arnosti DN. Mathematical modeling of gene expression: a guide for the perplexed biologist. Crit. Rev. Biochem. Mol. Biol. 2011;46(2): 137–151.

4. Bintu L, Buchler NE, García HG, Gerland U, Hwa T, Kondev J, Phillips R. Transcriptional regulation by the numbers: models. Current Opinion in Genetics & Development 2005;15: 116–124.

5. Bisswanger H. Encyme Kinetics: Principles and methods. 2nd ed. Weinheim: WILEY-VCH; 2008.

6. von Dassow G, Meir E, Munro E M, Odell GM. The segment polarity network is a robust developmental module. Nature. 2000;406: 188–192.

7. Frank TD, Carmody AM, Kholodenko BN. Versatility of Cooperative Transcriptional Activation: A Thermodynamical Modeling Analysis for Greater-Than-Additive and Less-Than-Aditive Effects. PLoS ONE. 2012;7(4).

8. Frank TD, Cavadas MAS, Nguyen LK, Cheong A. Non-linear Dynamics in Transcriptional Regulation: Biological Logic Gates. Nonlinear Dynamics in Biological Systems, SEMA SIMAI Springer Series. 2016;7: 43–62.

9. Gilman A, Arkin AP. Genetic “C1400”: Representation and Dynamical Models of Genetic Components and Networks. Annu. Rev. Genomics Hum. Genet. 2002;3: 341–369.

10. Junker JP, Peterson KA, Nishi Y, Mao J, McMahon AP, van Oudenaarden A. A Predictive Model of Bifunctional Transcription Factor Signaling during Embrionic Tissue Patterning. Developmental Cell. 2014;31: 448–460. May 23, 201820/21

11. Lai K, Robertson MJ, Schaffer DV. The Sonic Hedgehog Signaling System as a Bistable Genetic Switch. Biophysical Journal. 2004;86: 2748–2757.

12. Muller B, Basler K. The repressor and activator forms of Cubitus interruptus control Hedgehog target genes trough common generic Gli-binding sites. Development. 2000;127: 2999–3007.

13. Parker DS, White MA, Ramos AI, Cohen BA, Barolo S. The cis-Regulatory Logic of Hedgehog Gradient Responses: Key Roles for Gli Binding Affinity, Competition, and Cooperativity. Sci. Signal. 2011;4: 1–16.

14. Ptashne M, Gann A. Transcrition activation by recruitment. Nature. 1997;386(10): 569–577.

15. Ptashne M. Regulation of transcription: from lambda to eukaryotes. Trends in Biochemical Sciences. 2005;30(6): 275–279.

16. Ramos AI, Barolo S. Low-affinity transcription factor binding sites shape morphogen responses and enhancer evolution. Philos. Trans. R. Soc. Lond. B. Biol. Sci. 2013;368(1632): 20130017.

17. Rogers KW, Schier AF. Morphogen Gradients: From Generation to Interpretation, Annu. Rev. Cell Dev. Biol. 2011;27: 377–407.

18. Saha K, Schaffer DV. Signal dynamics in Sonic hedgehog tissue patterning. Development. 2006;133: 889–900.

19. Shea M, Ackers GK. The OR Control System of Bacteriophage Lambda. A Physical-Chemical Model for Gene Regulation. J. Mol. Biol. 1985;181: 211–230.

20. Tabata T, Takei Y. Morphogenes, their identification and regulation. Development. 2004;131: 703–712.

21. Torroja C, Gorfinkiel N, Guerrero I. Mechanisms of Hedgehog Gradient Formation and Interpretation. Developmental Neurobiology. 2005;64: 334–356.

22. Verbeni M, Sanchez O, Mollica E, Siegl-Cachedenier I, Carlenton A, Guerrero I, Ruiz i Altaba A, Soler J. Morphogenetic action through flux-limited spreading. Physics of Life Reviews. 2013;10: 457–475.

23. White MA, Parker DS, Barolo S, Cohen BA. A model of spatially restricted transcription in opposing gradients of activators and repressors. Mol. Syst. Biol. 2012;8 (614).

24. Zhou X, Su Z. tCal: transcriptional probability calculator using thermodynamic model. Bioinformatics. 2008;24(22): 2639–2640.

